# The BLENDS Method for Data Augmentation of 4-Dimensional Brain Images

**DOI:** 10.1101/2021.06.02.446748

**Authors:** Kevin P. Nguyen, Vyom Raval, Abu Minhajuddin, Thomas Carmody, Madhukar H. Trivedi, Richard B. Dewey, Albert A. Montillo

## Abstract

**Purpose:** Data augmentation improves the accuracy of deep learning models when training data is scarce by synthesizing additional samples. This work addresses the lack of validated augmentation methods specific for synthesizing anatomically realistic 4D (3D+time) images for neuroimaging, such as fMRI, by proposing a new augmentation method.

**Materials and Methods:** The proposed method, BLENDS, generates new nonlinear warp fields by combining intersubject coregistration maps, computed using symmetric normalization, through spatial blending. These new warp fields can be applied to existing 4D fMRI to create new augmented images. BLENDS is tested on two neuroimaging problems using de-identified datasets: 1) the prediction of antidepressant response from task-based fMRI in the EMBARC dataset (n = 163), and 2) the prediction of Parkinson’s Disease symptom trajectory from baseline resting-state fMRI regional homogeneity in the PPMI dataset (n = 43).

**Results:** BLENDS readily generates hundreds of new fMRI from existing images, with unique anatomical variations from the source images, that significantly improve prediction performance. For antidepressant response prediction, augmenting each original image once (2x the original training data) significantly increased prediction *R*^2^ from 0.055 to 0.098 *(p* < 1*e*^-6^), while at 10x augmentation *R*^2^ increased to 0.103. For the prediction of Parkinson’s Disease trajectory, 10x augmentation *R*^2^ increased from 0.294 to 0.548 *(p* < 1*e*^-6^).

**Conclusion:** Augmentation of fMRI through nonlinear transformations with BLENDS significantly improves the performance of deep learning models on clinically relevant predictive tasks. This method will help neuroimaging researchers overcome dataset size limitations and achieve more accurate predictive models.

## 1. INTRODUCTION

Data augmentation is an important tool for improving the performance of machine learning models, especially when limited data is available for training. Augmentation aims to simulate new data samples by introducing randomly generated variations to existing samples. Many applications of machine learning and deep learning to neuroimaging are limited by small dataset sizes, which could be readily addressed by augmentation. However, there are currently no augmentation methods to synthesize new, anatomically realistic functional magnetic resonance images (fMRI), complete with four-dimensional (3D + time) data.

The proposed method, Brain Library Enrichment through Nonlinear Deformation Synthesis (BLENDS), is the first that be applied to 4D MRI data including fMRI to generate new image timeseries. It is computationally efficient, requiring only 2 minutes on most hardware to generate each new image. It is also easily scalable and can generate up to hundreds of augmented images per original image by introducing brain morphological variations derived from large public repositories of MRI. Minimal user parameterization is required, making BLENDS straightforward to apply. BLENDS is shown to improve deep learning predictor accuracy in two use cases: prediction of antidepressant treatment outcome from task-based fMRI and Parkinson’s Disease total symptom trajectory prediction from resting-state fMRI. Beyond fMRI, the same method could also be applied to other neuroimaging modalities, including structural or diffusion MRI. Source code is available on a public repository and is ready for community use and extension.

Prior work in MRI augmentation has primarily focused on augmentation of T1-weighted structural MRI. Previously algorithms were developed that used ICA decomposition and random latent vectors to generate new T1-weighted images and improve a deep learning classifier of schizophrenic vs. healthy individuals (1, 2). Minimal work exists on methods for other brain MRI contrasts. Zhuang et al. proposed the synthesis of 3D task fMRI brain activation maps using a generative adversarial network (GAN) (3). However, we are the first to present a method to generate augmented 4D fMRI timeseries and demonstrate applications to multiple regression problems. This approach is compared to a previously reported algorithm in which augmented images by performing coregistrations with other, randomly selected images (4). A substantial improvement to the prediction of antidepressant outcomes was shown, however the method was limited by high computational cost and reliance on a two-step T1-based registration. This registration approach was limited to introducing only anatomical variations present in the sample at hand. The current algorithm, proposed herein, overcomes these limitations and demonstrates marked improvement in deep learning performance for two clinical problems while simultaneously reducing computation time by 15x and also generating out-of-sample augmentations.

## 2. METHODS

### 2.1. AUGMENTATION

A given brain’s morphology ***I*** can be nearly completely characterized by a warp field and a brain template. The brain template, such as the popular Montreal Neurological Institute 152-brain (MNI152) template, serves as an initial condition ***T***. A warp field ***W*** is a three-dimensional vector field which describes the nonlinear transformation of each point in the brain template to the given brain. For a specific point (*x, y, z*) in space, the transformation between ***T*** and ***I*** is modeled as:

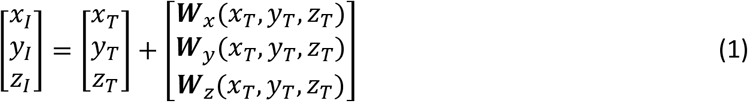

where ***W**_x_, **W**_y_, **W**_z_* are the x, y, and z components of the three-dimensional vector field. This warp field can be accurately computed at the voxel level through modern nonlinear coregistration tools such as ANTs and FNIRT (5, 6).

In the proposed method, a *set* of hundreds of warp fields are produced by coregistering T1-weighted images to the MNI152 template. ANTs Symmetric Normalization is used to perform an initial rough affine registration, followed by a multi-scale nonlinear registration (6). The resultant affine matrix for each image is decomposed into translation, rotation, shearing, and scaling components. The shearing and anisotropic scaling components are then composed with the nonlinear registration to obtain the warp field ***W*** which includes the non-rigid body and nonlinear transformation components. The rigid body translation and rotation components are excluded as they characterize the head position within the scanner bore, rather than characteristics of brain morphology. The generated warps are stored in set **Π** containing [***W***_1_, ***W***_2_,..., ***W***_*j*_] for *j* generated warps.

#### 2.1.1. Blended warp generation

Previous work showed that warping a given brain to a different, randomly selected brain through coregistration was effective for augmentation (4). However, this previous method could not introduce morphological variations beyond what exists in the dataset. Also, performing these coregistrations on-the-fly required ~30 minutes for every augmentation, which quickly becomes computationally costly when 100-1000 or more augmented images are desired. BLENDS eliminates this most costly step by generating new warps directly from the set of precomputed warps. Thus, an inordinate number of new warps can be rapidly created for the one-time fixed cost of creating the warpset. In many cases, a neuroimaging researcher may already have such a warp set on hand which may be repurposed for BLENDS. To generate new warps from the set, fully learning the distribution of warp fields, particularly at the voxel level, is computationally costly and unnecessary. Instead, a new warp can be created by spatially blending combinations of existing warps. With the assumption that a weighted combination of real anatomical warps yields an anatomically realistic warp, this approach allows the combinatorial generation of far more warps than are present in the set (Figure 1).

**Figure 1.**
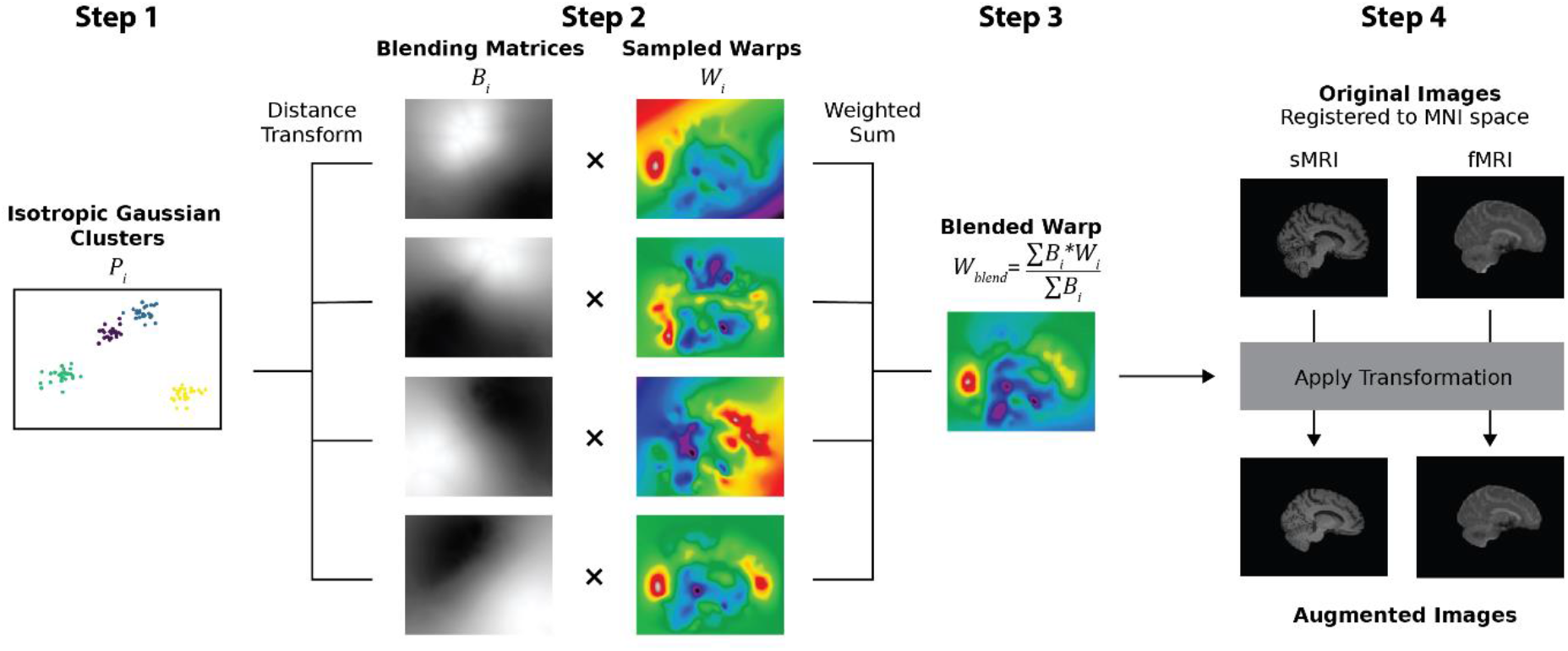
fMRI augmentation using BLENDS to combine 4 existing warp fields ***W**_i_* for *i* = 1, …, 4. Steps shown include: 1) Isotropic Gaussian point clusters in 3D space are generated for each of the 4 warp fields. 2) The distance transform is applied to each point cluster to smooth blending matrices ***B**_i_*. 3) The blending matrices are used to compute a weighted sum of the warp fields, resulting in a new blended warp ***W**_blend_*. 4) The blended warp is used to transform a pair of sMRI and fMRI images and produce a new, augmented sample. Colors in the warp fields indicate principal direction of local vector (red: sagittal, green: coronal, blue: axial).

For each augmentation operation, *n* real warps, ***W**_i_* for *i* = 1 … *n*, are drawn from the set **Π** and blended through a weighted sum. First, isotropic Gaussian point clusters *P_i_* are generated in 3D space for each warp, with random centroids within the image dimensions and standard deviation *σ.* Next, these discrete point clusters are converted into smooth, continuous 3D matrices ***B**_i_* through a distance transform:

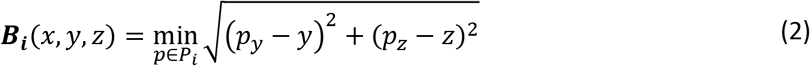

which computes, for each voxel (*x,y, z*) in ***B**_i_*, the Euclidean distance to the nearest point *p* in *P_i_*. The sagittal dimension *x* is omitted from the calculation to ensure left-right symmetry in the blended warp. Finally, the new, blended warp is computed as the sum of the real warps weighted by the blending matrices:

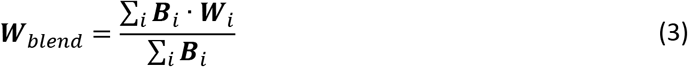

In the following applications, BLENDS was performed with the number of sampled real warps *n* = 4 and Gaussian point cluster *σ* = 1 (unitless). Point cluster coordinates were then min-max scaled to span the image dimensions (i.e. such that the range of coordinate values was (0, image size in voxels) for each dimension).

#### 2.1.2. Image preparation

The image to be augmented is the source image. In order to apply the newly generated warp ***W**_biend_* to the source image, the image must first be spatially normalized to the brain template, i.e. the initial condition. This is performed through a direct coregistration to an MNI152 EPI template. Registration to an EPI (i.e. fMRI template) has been shown to be more robust to EPI magnetic susceptibility artifacts and distortions than the conventional two-step T1-based normalization (EPI to T1 image, T1 image to template) (7, 8). This coregistration step is the most computationally expensive step of the augmentation procedure but can be completed in around 30 minutes with ANTs Symmetric Normalization (6). Importantly, this step need only be performed once per source image, irrespective of the number of desired augmented images.

Since the warps can be applied agnostically to any 3D or 4D image, the same generated warp can be used to transform multiple contrasts from the same individual such as fMRI, structural MRI, diffusion MRI, etc. In the case of a 4D fMRI timeseries, the warp is applied to each volume in the timeseries, assuming that the volumes have been realigned with a tool such as FSL MCFLIRT to correct for inter-volume motion (9).

### 2.2. APPLICATIONS

BLENDS is demonstrated on two distinct neuroimaging use cases involving common brain disorders. Both use de-identified, publicly available data containing fMRI and structural MRI (sMRI). Brief details are provided here with full dataset characteristics and MRI parameters available in the Supplemental Tables S1-4.

#### 2.2.1. Major depressive disorder

The objective in the first application is to predict future antidepressant response from brain activation measures of pre-treatment fMRI. Imaging data for 163 participants with major depressive disorder (MDD) are obtained from the Establishing Moderators and Biosignatures of Antidepressant Response in Clinical care (EMBARC) study (10). Demographics are shown in Supplemental Table S1. Functional imaging during a reward processing task and structural imaging were acquired at pretreatment baseline, before the participants underwent an 8-week course of the antidepressant sertraline (task paradigm and acquisition parameters are detailed in Supplemental Methods Section 1 and Supplemental Table S2). Depression severity was measured by trained clinical raters using the Hamilton Rating Scale for Depression 17-item score (HAMD) before and after treatment. Antidepressant response is defined as the difference in HAMD score.

The precomputed set of warps is generated by computing coregistrations between the MNI152 template and each of 290 structural (T1-weighted) images from EMBARC. BLENDS is applied to augment each fMRI twice (2x), five times (5x), and ten times (10x). Augmented fMRI are then preprocessed with a standard fMRI pipeline including head motion correction, spatial normalization, and nuisance regression (further details are provided in Supplemental Methods Sections 2 and 3). Brain activation maps are computed using a generalized linear model. A custom 200-region brain atlas was created from using the pyClusterROI tool, which performs spectral clustering normalized cuts on fMRI data (11). Mean regional values are computed using this atlas and passed as input features into a neural network to predict post-treatment change in HAMD score. The augmentation is performed on the “raw” 4D fMRI timeseries rather than the 3D brain activation maps. This both demonstrates the general applicability of BLENDS to 4D images and affords flexibility. In further applications, augmenting the original 4D images allows the user the ability to re-process and extract other measurements (e.g. functional connectivity, ReHo, fALFF, etc.) as needed for a targeted application.

#### 2.2.2. Parkinson’s Disease

For the second application, the goal is to predict Parkinson’s Disease (PD) total symptom trajectory from baseline resting-state fMRI. In this case, disease severity 12 months after baseline is to be predicted from the baseline regional homogeneity (ReHo) measures extracted from resting-state fMRI. From the Parkinson’s Progression Marker Initiative (PPMI) dataset, 43 PD participants are selected who had functional and structural imaging and who also had disease severity measured at 12 months after their baseline imaging time. Severity is measured by clinical raters using the total score from the Movement Disorder Society’s Unified Parkinson’s Disease Rating Scale (MDS-UPDRS), which was assessed with participants in the medicated state. See Supplemental Table S3 for demographics and Table S4 for acquisition parameters. The warp set contains 399 warps, computed from all available structural images from PPMI. As with the MDD application, BLENDS is applied at 2x, 5x, and 10x.

Augmented fMRI are preprocessed with the same fMRI pipeline (Supplemental Methods Sections 2 and 4). Regional homogeneity (ReHo), which measures the similarity in activity of each brain voxel with its 26-voxel neighborhood, is computed using the C-PAC software (v1.0.3) (12). ReHo reflects the synchronicity of a voxel with its immediate surrounding brain tissue, and abnormalities in ReHo have been observed in PD (13). The Schaefer 200-region atlas was used to compute the mean ReHo in each brain region. These values were used as inputs into the neural network, described in the next section, along with clinical covariates including age, sex, race, ethnicity, disease duration, and baseline MDS-UPDRS. The clinical covariates were not altered during the creation of each augmented sample.

### 2.3. NEURAL NETWORKS

Deep feed-forward, fully connected neural networks were constructed for both applications. Hyperparameter optimization was conducted using Bayesian Optimization with Hyperband (BOHB), a fully automated hyperparameter search algorithm unbiased to user expertise or hand-tuning (14). Full details of the hyperparameter optimization, model architectures, and hyperparameter ranges can be found in the Supplemental Methods Section 5, Figure S1, and Table S5.

Model performance was validated using nested K-fold cross-validation with 3 outer and 5 inner folds. For each outer fold, testing data was held aside and the remaining data was used to perform a random search over 100 hyperparameter configurations with 5-fold inner cross-validation. The best performing model from the random search was retrained on all inner data and evaluated on the testing data. This retraining and evaluation was repeated 100 times per outer fold, with different random weight initializations, for significance testing. No augmented data was included in validation or testing and data splits were grouped such that each augmented sample remained in the same fold as the original sample.

Neural networks were implemented in Tensorflow v1.12 and BOHB searches were performed with Ray Tune v1.2 with parallelization across 4 Nvidia Tesla P100 GPUs.

## 3. RESULTS

### 3.1. AUGMENTED IMAGES

Examples of augmented images generated with BLENDS are shown in Figure 2. A pair of structural and functional images from the same individual are augmented 5 times (same blended warp applied to both images) to introduce morphological variations throughout the brain. Comparing the augmented fMRI to the original image (Figure 2, 3^rd^ row) highlights areas where variations have been introduced: for example, the cerebellum size and the convexity of the frontal lobe have been altered to varying degrees. Due to the process of blending together multiple existing warps, these resultant augmented images are distinct from those in the original sample. Figure S2 illustrates examples of brain activation maps computed from original and augmented fMRI from the MDD dataset, showing that the variability is carried into subsequent measurements derived from fMRI. Additionally, each augmentation operation, including the generation of the blended warp and the transformation of the original image, required 2 minutes on a conventional workstation CPU (Xeon E5-2680) compared to 30 minutes required by our previous coregistration-based method, on the same hardware.

**Figure 2.**
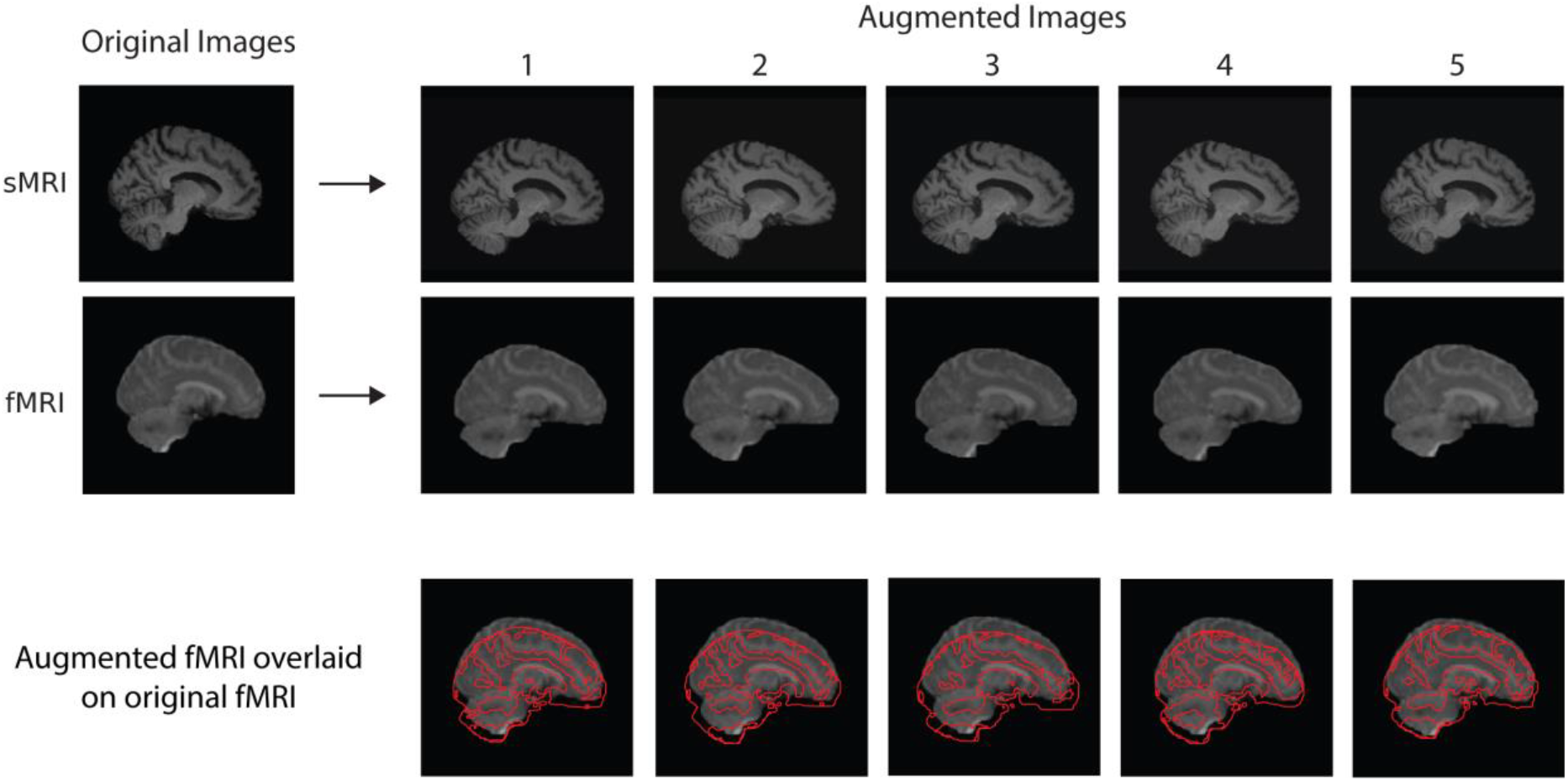
BLENDS applied to structural T1-weighted (sMRI, 1^st^ row) and functional MRI (fMRI, 2^nd^ row) from a Parkinson’s Disease patient to synthesize 5 new samples. Morphological variations are introduced in global brain shape as well as in neuroanatomical structures throughout the cerebellum and cerebrum. In the third row, an edge map of each augmented fMRI is overlaid over the original fMRI to illustrate the morphological differences.

### 3.2. MAJOR DEPRESSIVE DISORDER

Performance in predicting antidepressant response from task-based brain activation measures substantially increased with augmentation (Figure 3). At baseline before augmentation, *R*^2^ (mean over all outer cross-validation folds) was 0.055. This indicates that the model was able to explain about 5.5% of the variance in the change in depression symptoms 8 weeks after antidepressant treatment from the pre-treatment neuroimaging. Performance significantly increased with 2x augmentation to 0.098 *(p* < 1*e*^-6^), and it continued to increase with 5x (*R*^2^ 0.099) and 10x (*R*^2^ 0.103) augmentation. Thus with BLENDS augmentation the percent variance explained nearly doubled from 5.5% to 9.9%, a substantial and statistically significant improvement.

**Figure 3.**
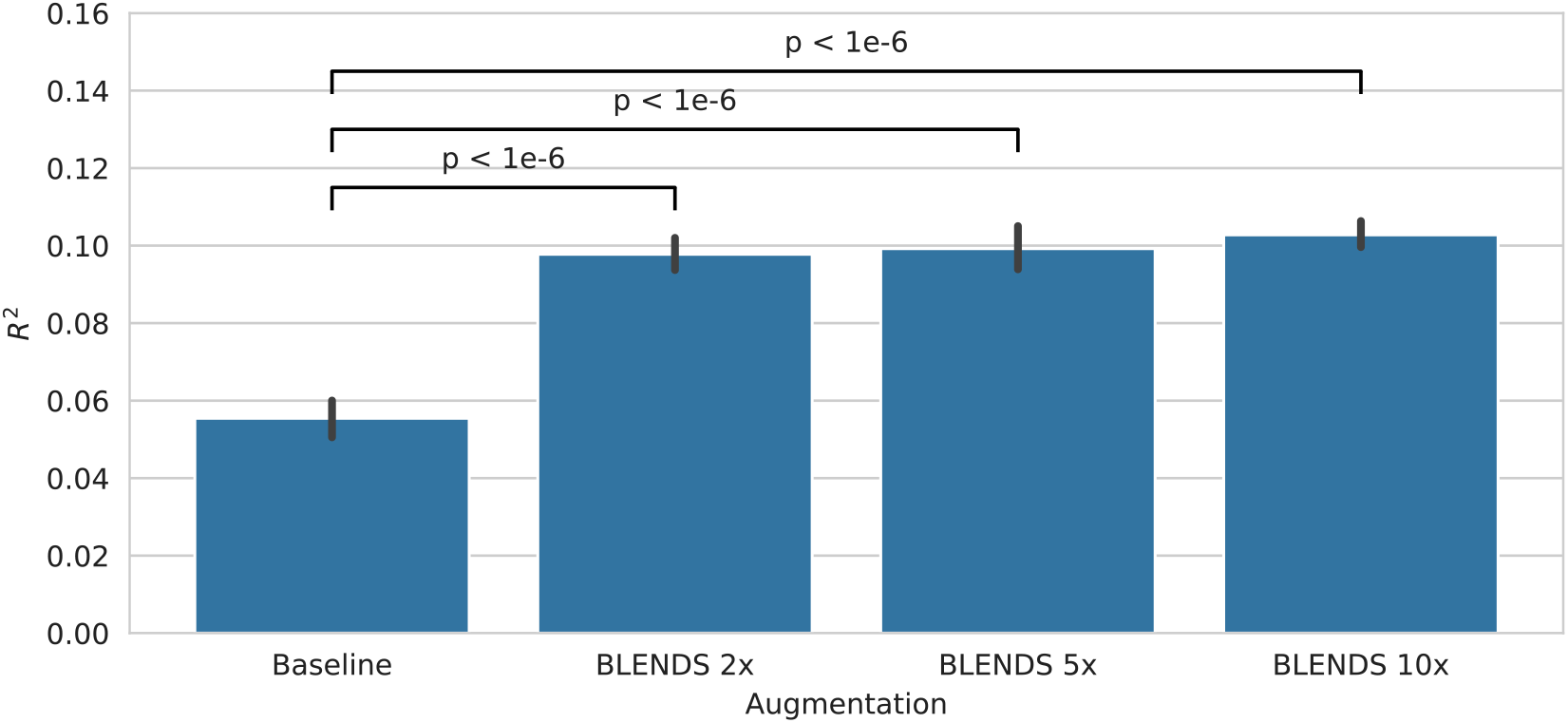
Performance (coefficient of determination, *R*^2^) in predicting antidepressant outcomes from pre-treatment task-based fMRI. Error bars indicate the 95% confidence interval, computed by retraining each model 100 times per cross-validation fold with different random weight initializations.

### 3.3. PARKINSON’S DISEASE

BLENDS also markedly improved the accuracy of predicting Parkinson’s Disease total MDS-UPDRS severity at 12 months after the baseline imaging scan using regional homogeneity measures (Figure 4). Before augmentation, *R*^2^ was 0.294, indicating that 29% of the variance in the total symptom level in the 12 months after baseline imaging was explained by the model. A significant increase to an *R*^2^ of 0.535 was seen with 5x augmentation *(p* < 1*e*^-6^). The trend continued with 10x augmentation *(R*^2^ 0.548). Thus again, with BLENDS augmentation increased from 29% to 55%, which is also a substantial and statistically significant improvement.

**Figure 4.**
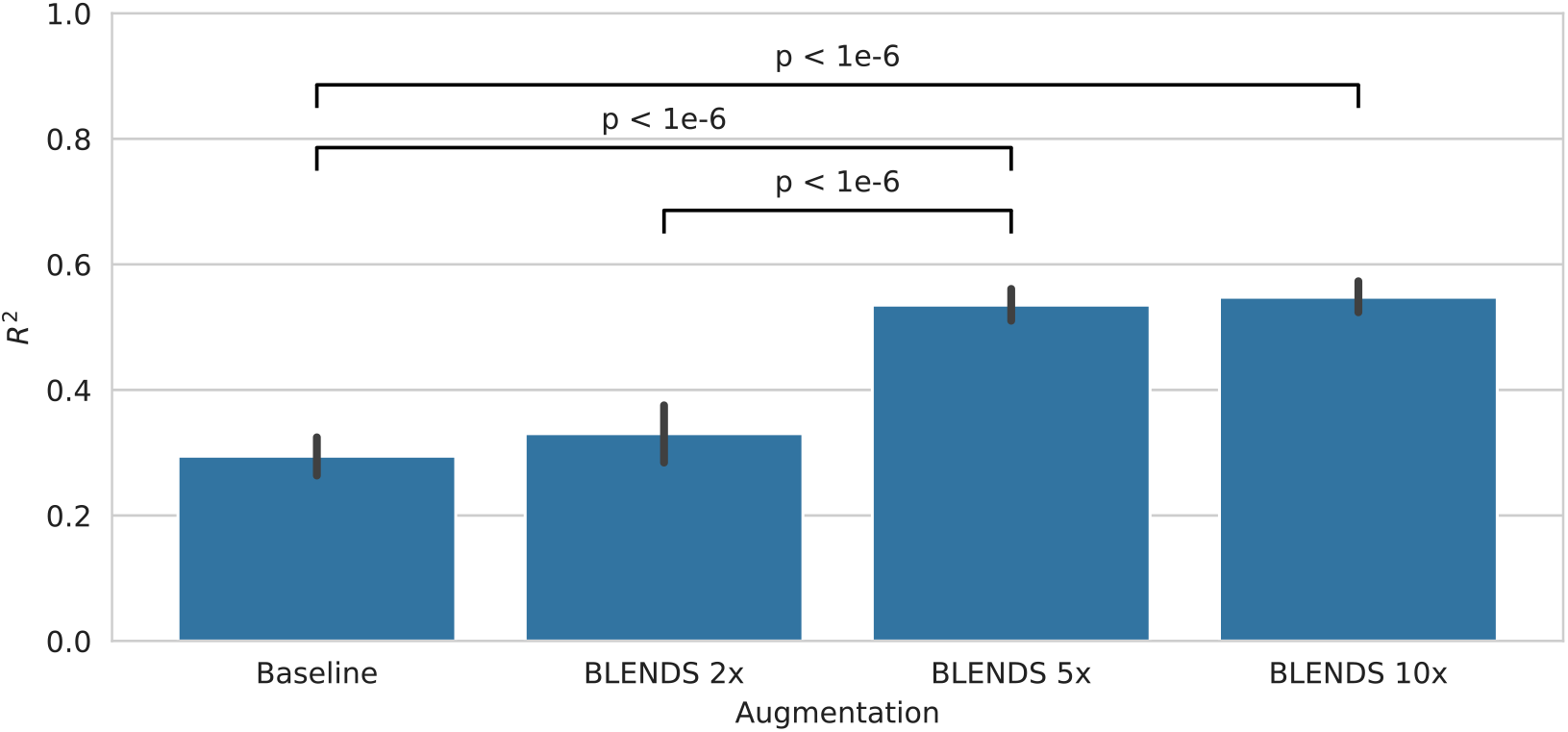
Performance (coefficient of determination, *R*^2^) in predicting future Parkinson’s Disease severity 1 year in the future from regional homogeneity derived from baseline resting-state fMRI. Error bars indicate the 95% confidence interval, computed by retraining each model 100 times per crossvalidation fold with different random weight initializations.

## 4. DISCUSSION

### 4.1. IMPACT ON PREDICTOR PERFORMANCE

For both clinical applications, the prediction of antidepressant response from task-based fMRI and the prediction of future Parkinson’s Disease severity from resting-state fMRI, BLENDS augmentation substantially improved neural network performance. In the antidepressant application, treatment outcome prediction performance improved by a relative 87% with 10x augmentation compared to baseline. For the prognosis of individuals with Parkinson’s Disease, BLENDS also demonstrated a large and significant performance increase from baseline. At 10x augmentation, performance was 86% greater relative to baseline. Of note, the relationship between augmentation amount and performance is not linear. Consequently, for a new problem, it may not be known *a priori* what amount of augmentation is required to achieve a desired performance level. However, a straightforward search over augmentation amount can be employed to identify the optimal condition.

Besides (4), there are few published fMRI augmentation methods for comparison. Zhuang et al. recently reported an approach using a generative adversarial network (GAN) to augment 3D brain activation maps derived from fMRI (3). They achieve up to a 4.6% accuracy increase (7.6% *relative* increase from baseline) in classifying cognitive states with a neural network. A main limitation of this approach is that it is constrained to 3D images; 4D GANs are computationally prohibitive to train on current hardware due to the sheer memory and compute required. Thisprevents usage on “raw” fMRI timeseries (3D + time) or other 4D data such as dynamic contrast-enhanced MRI or diffusion MRI (where the fourth dimension is direction). Similarly, Eslami et al. propose an extension to the Synthetic Minority Over-sampling Technique (SMOTE) using a similarity measure optimized for fMRI data (15). However, this method is also limited to generating fMRI derivatives (e.g. functional connectivity matrices) instead of full 4D images. Augmentation of the full fMRI timeseries is essential if subsequent analysis requires deriving scalar measures, such as functional connectivity or causal connectivity, or multiple complementary measures such as regional homogeneity (ReHo) and amplitude of low frequency fluctuations (ALFF). Instead of directly synthesizing images, BLENDS generates new warps that can be applied to any image type, allowing more flexibility than a GAN-based approach. An additional limitation to these existing methods is that they are restricted to classification tasks (i.e. with a discrete prediction target), which prevents application to the regression tasks investigated in this work. The GAN approach generates samples conditioned on the desired target label, and GANs conditioned on continuous labels do not yet exist. SMOTE is inherently limited to classification and while an extension to regression exists (SMOTER), it depends on a linearity assumption to interpolate new target values (16).

To demonstrate applicability to classification problems, BLENDS was also applied to the *diagnosis* of Parkinson’s Disease from resting-state functional connectivity (see Supplemental Methods Section 6). On this problem, BLENDS achieved a similar increase to classification accuracy as SMOTE, up to 10% compared to the baseline without augmentation (Figure S3). This highlights how BLENDS can achieve competitive performance boosting augmentation results for *both* regression and classification applications.

### 4.2. LIMITATIONS

There are two limitations of BLENDS that warrant discussion. The *first* is that BLENDS requires a precomputed set of warps, which can entail a large upfront computational cost. However, the combinatorial nature of sampling and blending warps means that a large range of morphological diversity can be generated from a relatively small warp set. For example, sampling 4 warps at a time from a set of 100 warps yields 4 million unique combinations, with further variations introduced through the random blending process. To mitigate this limitation, we will provide a large set of warps computed from over 1000 healthy brains to the community for ready download, facilitating immediate testing and application of BLENDS. For applications where the cohort may have aberrant morphology compared to healthy brains (e.g. Parkinson’s Disease), we recommend that researchers compute their own warp set. In many cases, researchers already have such a set of study specific warps computed, as coregistration of brains to an atlas is a common step in many preprocessing pipelines. The *second* limitation is that BLENDS may not be applicable to situations where random warping may perturb biological information of interest in images, including cases where morphology is directly associated with the prediction target.

However, this limitation is inherent to any data augmentation method that relies on random transformations. In the case of BLENDS, this could be partially mitigated by using stratified or separate sets for each class, as was done with the Parkinson’s Disease diagnosis application.

### 4.3. CONCLUSION

These results establish clear evidence to augment neuroimaging data for deep learning when the data is limited. BLENDS demonstrated a substantial performance benefit when applied to two distinct neuroimaging problems encompassing psychiatric and neurological disease. Main advantages to the method include its 1) flexibility in synthesizing full 4D images, 2) computational speed, and 3) ability to produce out-of-sample variations through warp blending. These results support the use of BLENDS for task-based and resting-state fMRI, though future work could investigate applications to other MRI contrasts, including diffusion or dynamic contrast enhanced MRI, where the four-dimensional data is costly and could benefit from augmentation. Consequently, BLENDS is anticipated to be of general interest to the neuroimaging community and especially to researchers looking to improve performance of deep learning on their existing data.

## ACKNOWLEDGEMENTS

We thank Drs. Andrew Jamieson, Jeon Lee, and Jian Zhou for their feedback during the writing of this manuscript.

Data used in the preparation of this article were obtained from the Parkinson’s Progression Markers Initiative (PPMI) database (www.ppmi-info.org/data). For up-to-date information on the study, visit www.ppmi-info.org. PPMI – a public-private partnership – is funded by the Michael J. Fox Foundation for Parkinson’s Research and funding partners, including Abbvie, Allergan, Amathus Therapeutics, Avid Radiopharmaceuticals, Biogen, BioLegend, Bristol-Myers Squibb, Celgene, Denali, GE Healthcare, Genentech, GlaxoSmithKline, Golub Capital, Handl Therapeutics, Insitro, Janssen Neuroscience, Lilly, Lundbeck, Merck, Meso Scale Discovery, Pfizer, Piramal, Prevail Therapeutics, Roche, Sanofi Genzyme, Servier, Takeda, Teva, UCB, Verily, and Voyager Therapeutics.

## CODE AVAILABILITY

To facilitate reuse and extension, the source code for our method can be found at https://github.com/DeepLearningForPrecisionHealthLab/BLENDS.

## REFERENCES

1. Castro E, Ulloa A, Plis SM, Turner JA, Calhoun VD. Generation of synthetic structural magnetic resonance images for deep learning pre-training. In: 2015 IEEE 12th International Symposium on Biomedical Imaging (ISBI). IEEE, 2015; 1057–1060.

2. Ulloa A, Plis S, Erhardt E, Calhoun V. Synthetic structural magnetic resonance image generator improves deep learning prediction of schizophrenia. In: 2015 IEEE 25th International Workshop on Machine Learning for Signal Processing (MLSP). [S.l.]: IEEE, 2015; 1–6.

3. Zhuang P, Schwing AG, Koyejo O. FMRI Data Augmentation Via Synthesis. In: IEEE International Symposium on Biomedical Imaging (ISBI). [Piscataway, New Jersey]: IEEE, 2019; 1783–1787.

4. Nguyen KP, Chin Fatt C, Treacher A, Mellema C, Trivedi MH, Montillo A. Anatomically-informed data augmentation for functional MRI with applications to deep learning. In: Landman BA, Išgum I, eds. Medical Imaging: Image Processing. SPIE, 2020; 28–33.

5. Andersson J, Jenkinson M, Smith S, 2007. Non-linear registration, aka Spatial normalisation: FMRIB Technical Report TR07JA2. Oxford.

6. Avants BB, Tustison NJ, Song G, Cook PA, Klein A, Gee JC. A Reproducible Evaluation of ANTs Similarity Metric Performance in Brain Image Registration. Neuroimage 2010;54(3):2033–2044. doi:10.1016/j.neuroimage.2010.09.025.

7. Calhoun VD, Wager TD, Krishnan A, et al. The impact of T1 versus EPI spatial normalization templates for fMRI data analyses. Hum Brain Mapp 2017;38(11):5331–5342. doi:10.1002/hbm.23737.

8. Dohmatob E, Varoquaux G, Thirion B. Inter-subject Registration of Functional Images: Do We Need Anatomical Images? Front Neurosci 2018;12:64. doi:10.3389/fnins.2018.00064.

9. Jenkinson M, Bannister P, Brady M, Smith S. Improved Optimization for the Robust and Accurate Linear Registration and Motion Correction of Brain Images. Neuroimage 2002;17(2):825–841. doi:10.1016/S1053-8119(02)91132-8.

10. Trivedi MH, McGrath PJ, Fava M, et al. Establishing moderators and biosignatures of antidepressant response in clinical care (EMBARC): Rationale and design. J Psychiatr Res 2016;78:11–23. doi:10.1016/j.jpsychires.2016.03.001.

11. Craddock RC, James GA, Holtzheimer PE, Hu XP, Mayberg HS. A whole brain fMRI atlas generated via spatially constrained spectral clustering. Hum Brain Mapp 2012;33(8):1914–1928. doi:10.1002/hbm.21333.

12. Craddock C, Sikka S, Cheung B, et al. Towards Automated Analysis of Connectomes: The Configurable Pipeline for the Analysis of Connectomes (C-PAC). Front Neuroinform 2013;7. doi:10.3389/conf.fninf.2013.09.00042.

13. Hu J, Xiao C, Gong D, Qiu C, Liu W, Zhang W. Regional homogeneity analysis of major Parkinson’s disease subtypes based on functional magnetic resonance imaging. Neuroscience Letters 2019;706:81–87. doi:10.1016/j.neulet.2019.05.013.

14. Falkner S, Klein A, Hutter F. BOHB: Robust and Efficient Hyperparameter Optimization at Scale. In: Jennifer Dy, Andreas Krause, eds. Proceedings of the 35th International Conference on Machine Learning. Stockholmsmässan, Stockholm Sweden: PMLR, 2018; 1437–1446.

15. Eslami T, Mirjalili V, Fong A, Laird AR, Saeed F. ASD-DiagNet: A Hybrid Learning Approach for Detection of Autism Spectrum Disorder Using fMRI Data. Front Neuroinform 2019;13:70. doi:10.3389/fninf.2019.00070.

16. Torgo L, Ribeiro RP, Pfahringer B, Branco P. SMOTE for Regression. In: Correia LM, Reis LP, Cascalho J, eds. Progress in artificial intelligence: 16th Portuguese Conference on Artificial Intelligence, EPIA 2013, Angra do Heroísmo, Azores, Portugal, September 9-12, 2013, proceedings / Luís Correia, Luís Paulo Reis, José Cascalho (eds.). Heidelberg: Springer, 2013; 378–389.

