## Supplemental Material for "The BLENDS Method for Data Augmentation of 4-Dimensional Brain Images"

#### SUPPLEMENTAL TABLES AND FIGURES

**Table S1**. EMBARC dataset demographics.

| | |
|---|---|
| Sample size | 163 |
| Mean age | 38.8 ± 13.6 years |
| Female | 64.4% |
| Baseline HAMD | 18.6 ± 4.2 |
| Mean HAMD change | 7.5 ± 7.0 |

**Table S2**. MRI acquisition parameters for each EMBARC study site.

| | **Columbia University** | **Massachusetts General Hospital** | **University of Michigan** | **UT Southwestern Medical Center** |
|---|---|---|---|---|
| Scanner | General Electric Signa HDx 3T | Siemens TrioTim 3T | Philips Achieva 3T | Philips Ingenia 3T |
| Structural MRI | | | | |
| Sequence | FSPGR | MPRAGE | TFE | MPRAGE |
| TR/TI/TE | 6.0ms/900ms/2.4ms | 2300ms/900ms/2.54ms | 8.2ms/1100ms/3.7 ms | 2100ms/1100ms/3.7ms |
| Flip angle | 9° | 9° | 12° | 12° |
| Dimensions | 256 x 256 x 174 | 256 x 256 x 176 | 256 x 256 x 178 | 256 x 256 x 178 |
| Voxel size | 1 x 1 x 1 mm | 1 x 1 x 1 mm | 1 x 1 x 1 mm | 1 x 1 x 1 mm |
| Functional MRI | | | | |
| Sequence | GE-EPI | GE-EPI | GE-EPI | GE-EPI |
| TR/TE | 2000ms/28ms | 2000ms/28ms | 2000ms/28ms | 2000ms/28ms |
| Flip angle | 90° | 90° | 90° | 90° |
| Dimensions | 64 x 64 x 39 | 64 x 64 x 39 | 64 x 64 x 39 | 64 x 64 x 39 |
| Voxel size | 3.2 x 3.2 x 3.1 mm | 3.2 x 3.2 x 3.1 mm | 3.2 x 3.2 x 3.1 mm | 3.2 x 3.2 x 3.1 mm |
| Dummy scans | 5 | 5 | 5 | 5 |
| Number of volumes | 240 | 240 | 240 | 240 |
| Total acquisition time | 480 s | 480 s | 480 s | 480 s |

**Table S3**. PPMI dataset demographics

| | |
|---|---|
| Sample size | 43 |
| Mean age | 63.6 ± 10.3 years |
| Female | 30.3% |
| Baseline MDS-UPDRS | 37.2 ± 16.7 |
| Disease duration | 103 ± 525 days |

**Table S4**. PPMI MRI acquisition parameters

| | |
|---|---|
| Scanner | Siemens TrioTim 3T |
| Structural MRI | |
| Sequence | MPRAGE |
| TR/TI/TE | 2300ms/900ms/2.98ms |
| Flip angle | 9° |
| Dimensions | 240 x 256 x 176 |
| Voxel size | 1 x 1 x 1 mm |
| Functional MRI | |
| Sequence | 2D-EPI |
| TR/TE | 2400ms/25ms |
| Flip angle | 80° |
| Dimensions | 68 x 66 x 33 |
| Voxel size | 3.3 x 3.3 x 3.3 mm |
| Number of volumes | 210 |
| Total acquisition time | 504 s |

**Table S5.** Hyperparameter search ranges

| Hyperparameter | Sampling Distribution |
|---|---|
| Deep feed forward neural network layer hyperparameters | |
| Number of fully-connected hidden layers, $L$ | Uniform(min=1, max=6) |
| Number of neurons in the first hidden layer, $N_1$ | Uniform(min=64, max=512) |
| Weight L1 regularization strength, $\lambda_{1,i}$ | Uniform(min=0.05, max=0.9) |
| Weight L2 regularization strength, $\lambda_{2,i}$ | Uniform(min=0.05, max=0.9) |
| Taper rate (% decrease in hidden layer size from previous layer), $t$ | Uniform(min=0.05, max=0.5) |
| Batch normalization, $b_i$ | [True, False] |
| Dropout rate, $r_i$ | Uniform(min=0.2, max=0.8) |
| Activation function, $a(\cdot)$ | [ReLU, LeakyReLU, ELU, PReLU] |
| Nadam Optimizer hyperparameters | |
| Learning rate | Uniform(min=0.001, max=0.030) |
| Momentum, $\beta_1$ | Uniform(min=0.3, max=0.9) |
| Momentum, $\beta_2$ | Uniform(min=0.6, max=0.9) |

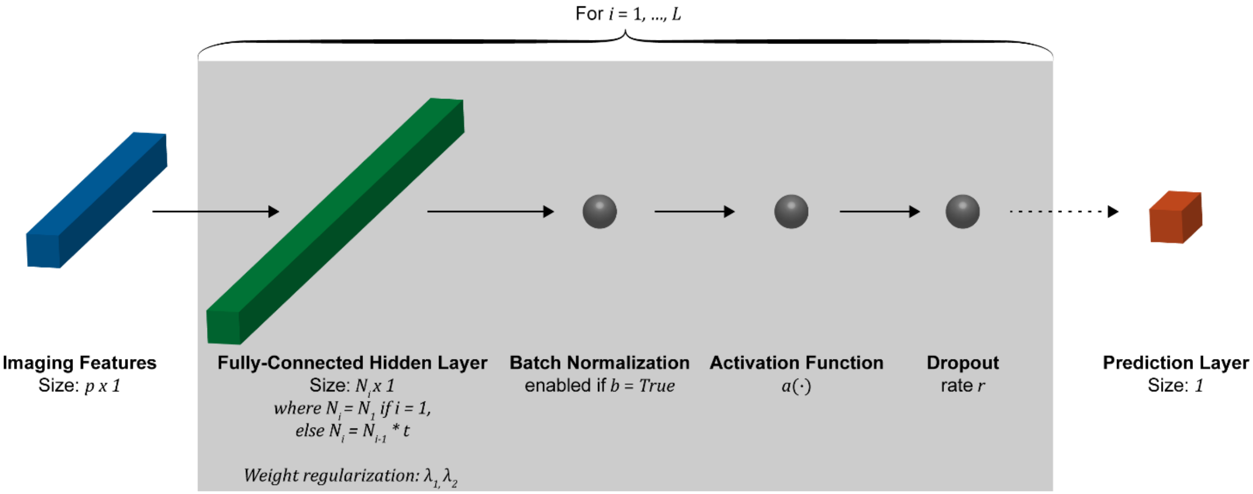


**Figure S1**. Deep feed-forward fully connected neural network architecture. Hyperparameters including number of hidden layers $L$, hidden layer size $N_i$, regularization $\lambda_1, \lambda_2$, batch normalization $b$, activation function $a(\cdot)$, and dropout rate $r$ were optimized using Bayesian Optimization with Hyperband (see Table S5). The size of each hidden layer $N_i$ was determined using the size of the previous hidden layer $N_{i-1}$ and the taper rate hyperparameter $t$.

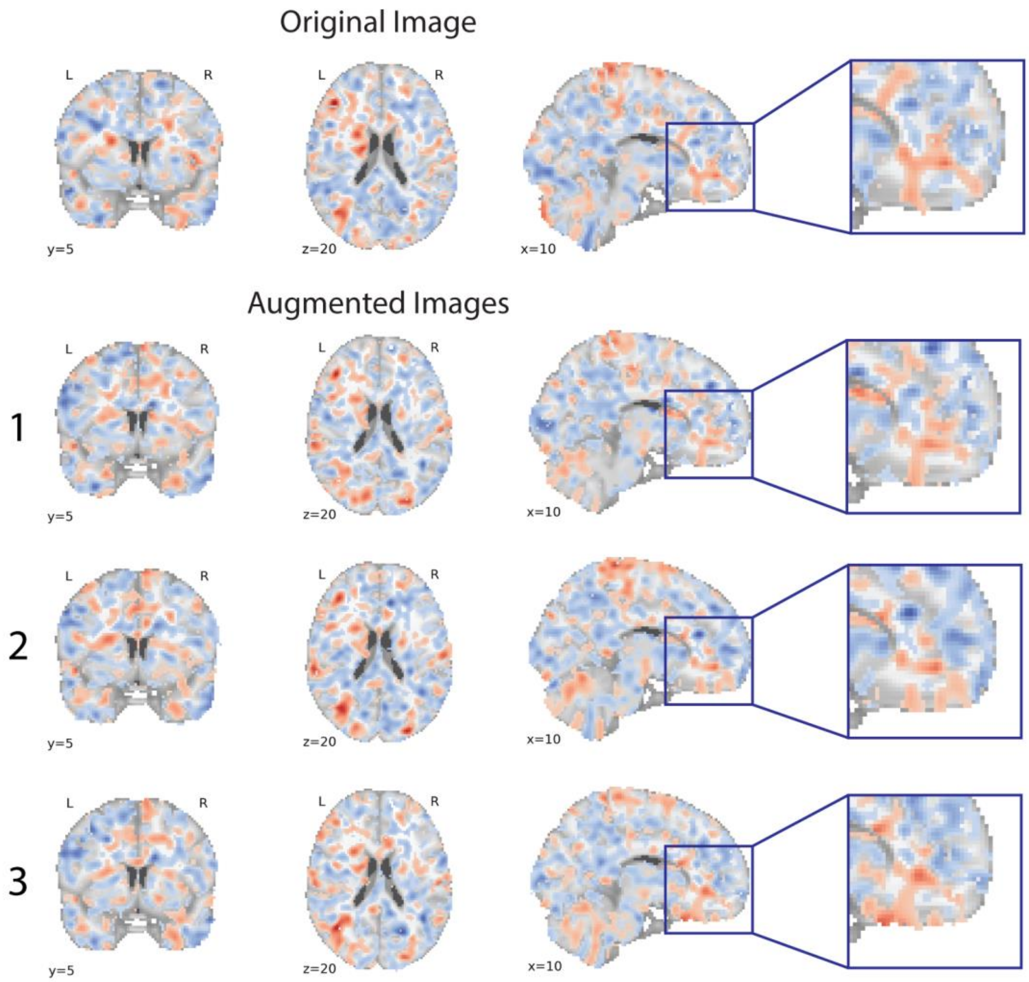


**Figure S2**. Brain activation maps computed from the original and three augmented fMRI of a participant from the major depressive disorder dataset. Red indicates areas of higher activity during reward stimuli and blue indicates areas of lower activity. Maps have been spatially normalized to the MNI152 template to control for global brain shape differences while highlighting local differences among the brain activation maps. An area of the prefrontal cortex has been magnified to show how augmentation has introduced variation in activation patterns.

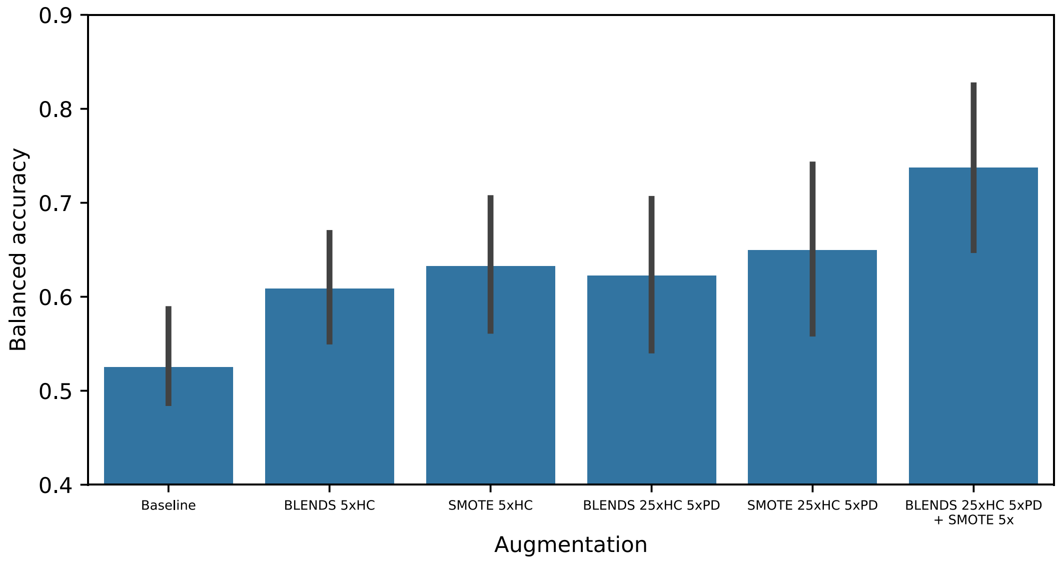


**Figure S3**. While BLENDS was designed for regression, even for classification of Parkinson's Disease (PD) vs. healthy control (HC) from resting-state functional connectivity, BLENDS provides an increase in performance similar to SMOTE. At baseline, severe class imbalance (5 PD: 1 HC) results in balanced accuracy of 52.5%. Correcting the class imbalance by augmenting the HC class 5x with either BLENDS or SMOTE improves accuracy to 60.9% and 63.3% respectively. Further augmenting the HC class 25x and PD class 5x increases accuracy to 62.3% and 63.3% respectively. Finally, augmenting the HC class 25x and PD class 5x with BLENDS, followed by further 5x augmentation with SMOTE (total augmentation was 125x for HC and 25x for PD), increases accuracy to 73.8%. Hence, the BLENDS and SMOTE methods are compatible and can be used in combination for classification problems. Error bars in this figure indicate the 95% confidence interval computed from 10 cross-validation folds.

### SUPPLEMENTAL METHODS

#### 1. EMBARC Reward Task Paradigm

The EMBARC major depressive disorder study acquired task-based fMRI using a number guessing task design to stimulate the reward processing areas of the brain. The task consists of 24 trials, divided into “possible win” and “possible loss” trials. During the “response phase” at the beginning of each trial, the participant guesses whether the upcoming number is greater or less than 5, with possible values ranging from 1-9. After pressing a button corresponding to their guess, the “anticipation phase” begins and they are informed about the trial type. In a possible win trial, the participant wins $1 for a correct guess and loses nothing for a wrong guess. In a possible loss trial, they win nothing for a correct guess and lose $0.50 for a wrong guess. The number is revealed in the “outcome phase” and the participant receives visual feedback about their monetary reward or penalty. A “baseline phase” occurs between each trial.

#### 2. Functional MRI Preprocessing Pipeline

All fMRI from both EMBARC and PPMI datasets were preprocessed through the following image processing pipeline. First, FSL MCFLIRT was used to correct inter-volume head motion and estimate affine head motion parameters (1). Brain extraction was performed by applying FSL BET and AFNI 3dAutomask and keeping the intersection of the two masks, an approach developed by the fMRIPrep pipeline (2–4). To spatially normalize to the MNI152 EPI template, the mean volume was coregistered to the template using ANTs Symmetric Normalization (5).

After this step of the pipeline, BLENDS was applied to augment the images. The augmented images were then smoothed with a 4mm FWHM Gaussian kernel and nuisance regression was performed to remove effects of head motion and physiological artifacts. Nuisance regressors included the 6 affine head motion parameters and their first derivatives in time and their squares, motion-related regressors computed by ICA-AROMA (6), and mean white matter and cerebrospinal fluid signals.

#### 3. EMBARC Feature Extraction

To compute brain activation maps from EMBARC fMRI, subject-level generalized linear models (GLMs) were fitted to each preprocessed image using SPM12. The GLM design matrix was based on well-validated previous analyses of this dataset (7, 8). Regressors were created for the response, anticipation, outcome, and baseline phases of the task. Parametrically modulated regressors were added to represent *reward expectation* and *prediction error*. The reward expectation regressor was set to the expected value of the monetary outcome of the two trial types, during the anticipation phase. For “possible win” trials, it had a value of +0.5 during the anticipation phase, while for “possible loss” trials, it had a value of -0.25 during the anticipation phase. The prediction error regressor was set to the difference between the monetary outcome of each trial and the expected value, during the outcome phase. For “possible win” trials, it had a value of +0.5 for correct guesses and -0.5 for wrong guesses. For “possible loss” trials, it had a value of +0.25 for correct guesses and -0.25 for wrong guesses.

The $time \times regressor$ design matrix $\boldsymbol{X}$ for the GLM comprised these 6 regressors and the GLM was defined as

$$\boldsymbol{Y} = \boldsymbol{X\beta} + \boldsymbol{\epsilon}$$

where $\boldsymbol{Y}$ is the observed $time \times voxels$ data matrix containing the voxel timeseries from the fMRI, $\boldsymbol{\beta}$ is the $regressors \times voxels$ matrix containing the fitted coefficients, and $\boldsymbol{\epsilon}$ is the $time \times voxels$ matrix containing residuals. The contrasts of interest include: 1) *anticipation* defined as $\boldsymbol{\beta}_{anticipation} - \boldsymbol{\beta}_{baseline}$, 2) *reward expectation* defined as $\boldsymbol{\beta}_{reward\ expectation}$, and 3) *prediction error* defined as $\boldsymbol{\beta}_{prediction\ error}$. A custom 200-region brain atlas was generated by applying the pyClusterROI spectral clustering tool to resting-state fMRI from EMBARC (9). This atlas was used to compute mean regional values for each contrast, and these values were used as inputs into the neural network.

#### 4. PPMI Feature Extraction

On the second dataset, PPMI, regional homogeneity (ReHo) was computed from the preprocessed fMRI to use as neural network inputs. ReHo was measured by computing Kendall's coefficient of concordance between each voxel and each of its 26 voxel neighbors (9) using the software C-PAC v1.0.3. Finally, the Schaefer 200-region brain atlas, similar in granularity to the custom atlas generated for EMBARC, was used to compute mean regional ReHo values, which were used as neural network inputs (10).

#### 5. Neural Network Hyperparameter Optimization

Bayesian optimization with hyperband (BOHB) was conducted to optimize the hyperparameters of the feed-forward fully-connected neural networks (Figure S1). BOHB automatically and efficiently searches a high-dimensional hyperparameter space and identifies a high-performing hyperparameter configuration in an unbiased manner (11). BOHB was implemented using the Ray Tune library in Python, and the hyperparameter ranges and distributions searched are enumerated in Table S5.

The BOHB approach was combined with nested K-fold cross-validation to select high-performing hyperparameter configurations and to evaluate them on held-out testing data. The data was first split into 3 outer folds, with stratification (by treatment outcome score for the antidepressant response application and 1-year severity score for the Parkinson's Disease application) to ensure similar distributions of labels for each fold. For each of these splits, the test fold was set aside and the remaining data was used for hyperparameter optimization. The hyperparameter optimization evaluated 100 distinct model configurations over the distributions detailed in Table S5. Each model was trained and evaluated with an inner 5-fold cross-validation. Augmented data was added to the training data after performing the cross-validation splits. The best performing model was selected based on lowest root mean squared error over all inner folds. This model was retrained on all inner fold data, and final performance was evaluated on the testing fold. To generate a distribution for significance testing, the final training and evaluation was performed 100 times with different random weight initializations. Finally, the entire process including BOHB and final evaluation was repeated for each of the 3 outer folds. The mean performance over all outer folds (300 total data points per augmentation condition) is reported in the main text.

Models were trained with a loss function that maximized the regression coefficient of determination $R^2$:

$$L = \left(1 - R^2(y, \hat{y})\right)$$

where $y$ and $\hat{y}$ are the true and predicted labels.

#### 6. Parkinson's Disease Diagnosis

As a complement to the two regression problems described in the main text, BLENDS was also applied to the diagnosis of Parkinson's Disease from resting-state functional connectivity. While the prognosis application in the main text aimed to predict future disease severity from baseline fMRI, here the goal is to classify Parkinson's Disease (PD) and healthy control (HC) participants. This application is motivated by the need for an automated tool to inform diagnoses for clinicians. Imaging data for 102 PD patients and 22 HC participants were obtained from the Parkinson's Progression Marker Initiative (PPMI) dataset. Demographics for this dataset are shown in Table S3 while acquisition parameters are detailed in Table S4. Resting-state functional and structural images were selected from the earliest timepoint available for each individual.

Separate precomputed warp sets were created for PD and HC to control for potential morphological differences in diseased vs. non-diseased brains. The PD pool contained 399 warps and the HC pool contained 183 warps computed from structural (T1-weighted) images. Because this dataset suffers both low sample size and class imbalance, augmentation strategies were chosen to correct for both limitations. BLENDS was first used to augment HC fMRI by a factor of 5, correcting for the 5:1 class imbalance between PD and HC. Next, HC was augmented 25x while PD was augmented 5x, maintaining the class balance while increasing overall dataset size by an additional factor of 5.

Augmented fMRI were preprocessed with the same fMRI pipeline as the other two applications. Timeseries were extracted for regions defined by the Schaefer 200-region atlas and functional connectivity between each pair of regions was computed using Pearson's correlation. Functional connectivity values were fed as input features into the neural network to classify PD vs HC.

BLENDS was compared to SMOTE, an existing method for data augmentation for classification problems which generates new samples through random linear interpolations between neighboring real samples (12). A modified SMOTE algorithm was tested, which was developed specifically for functional connectivity and employs the Extended Frobenius Norm to select nearest neighbor samples based on similarity of fMRI timeseries (13). In contrast to BLENDS which generates images, SMOTE operates directly on the functional connectivity features. Like BLENDS, SMOTE was applied first to augment HC 5x, then HC 25x and PD 5x.

Classification was performed by a feed-forward neural network, optimized using BOHB in an identical approach to the regression applications. Data was partitioned into 20% test set and 80% training set, and an inner 10-fold cross-validation was used during hyperparameter optimization. This was repeated 10 times with unique random train/test splits to generate 10 measurements of test performance per augmentation condition. Results are shown in Figure S3 and discussed in section 4.1.

### REFERENCES

1. Jenkinson M, Bannister P, Brady M, Smith S. Improved Optimization for the Robust and Accurate Linear Registration and Motion Correction of Brain Images. Neuroimage 2002;17(2):825–841. doi:10.1016/S1053-8119(02)91132-8.
2. Esteban O, Markiewicz CJ, Blair RW, et al. fMRIPrep: a robust preprocessing pipeline for functional MRI. Nat Methods 2019;16(1):111–116. doi:10.1038/s41592-018-0235-4.
3. Smith SM. Fast robust automated brain extraction. Hum Brain Mapp 2002;17(3):143–155. doi:10.1002/hbm.10062.
4. Cox RW. AFNI: Software for Analysis and Visualization of Functional Magnetic Resonance Neuroimages. Computers and Biomedical Research 1996;29(3):162–173. doi:10.1006/cbmr.1996.0014.
5. Avants BB, Tustison NJ, Song G, Cook PA, Klein A, Gee JC. A Reproducible Evaluation of ANTs Similarity Metric Performance in Brain Image Registration. Neuroimage 2010;54(3):2033–2044. doi:10.1016/j.neuroimage.2010.09.025.
6. Pruim RHR, Mennes M, van Rooij D, Llera A, Buitelaar JK, Beckmann CF. ICA-AROMA: A robust ICA-based strategy for removing motion artifacts from fMRI data. Neuroimage 2015;112:267–277. doi:10.1016/j.neuroimage.2015.02.064.
7. Greenberg T, Chase HW, Almeida JR, et al. Moderation of the Relationship Between Reward Expectancy and Prediction Error-Related Ventral Striatal Reactivity by Anhedonia in Unmedicated Major Depressive Disorder: Findings From the EMBARC Study. Am J Psychiatry 2015;172(9):881–891. doi:10.1176/appi.ajp.2015.14050594.
8. Greenberg T, Fournier JC, Stiffler R, et al. Reward related ventral striatal activity and differential response to sertraline versus placebo in depressed individuals. Mol Psychiatry 2019;25(7):1526–1536. doi:10.1038/s41380-019-0490-5.
9. Craddock C, Sikka S, Cheung B, et al. Towards Automated Analysis of Connectomes: The Configurable Pipeline for the Analysis of Connectomes (C-PAC). Front Neuroinform 2013;7. doi:10.3389/conf.fninf.2013.09.00042.
10. Schaefer A, Kong R, Gordon EM, et al. Local-Global Parcellation of the Human Cerebral Cortex from Intrinsic Functional Connectivity MRI. Cereb Cortex 2018;28(9):3095–3114. doi:10.1093/cercor/bhx179.
11. Falkner S, Klein A, Hutter F. BOHB: Robust and Efficient Hyperparameter Optimization at Scale. In: Jennifer Dy, Andreas Krause, eds. Proceedings of the 35th International Conference on Machine Learning. Stockholmsmässan, Stockholm Sweden: PMLR, 2018; 1437–1446.
12. Chawla NV, Bowyer KW, Hall LO, Kegelmeyer WP. SMOTE: Synthetic Minority Over-sampling Technique. Journal of Artificial Intelligence Research 2002;16:321–357. doi:10.1613/jair.953.
13. Eslami T, Mirjalili V, Fong A, Laird AR, Saeed F. ASD-DiagNet: A Hybrid Learning Approach for Detection of Autism Spectrum Disorder Using fMRI Data. Front Neuroinform 2019;13:70. doi:10.3389/fninf.2019.00070.
